# Comparative Analysis of Machine Learning Libraries for Neural Networks: A Benchmarking Study

**DOI:** 10.1101/2025.02.02.635632

**Authors:** Aws Ismail Abu Eid, Salameh A. Mjlae, Suzie Yaseen Rababa’h, Ahmed Hammad, Mahmoud Ali Ibrahim Rababah

## Abstract

This comprehensive benchmarking study explores the performance of three prominent machine learning libraries: PyTorch, Keras with TensorFlow backend, and Scikit-learn with the same criteria, software, and hardware. The evaluation encompasses two diverse datasets, “student performance” and “College Attending Plan Classification,” supported by Kaggle platforms utilizing feedforward neural networks (FNNs) as the modeling technique. The findings reveal that PyTorch and Keras with TensorFlow backend excel on the “College Attending Plan Classification” dataset, with PyTorch achieving impeccable precision, Recall, and F1-score for both classes. While Scikit-learn demonstrates commendable performance, it trails behind these libraries in this context. On the “Student Performance” dataset, all three libraries deliver comparable results, with Scikit-learn exhibiting the lowest accuracy at 16%. Keras with TensorFlow backend and PyTorch attain accuracy rates of 23%, respectively. Moreover, this study offers valuable insights into each library’s unique strengths and weaknesses when confronted with diverse dataset types. PyTorch emerges as the go-to choice for demanding tasks requiring high performance, while Scikit-learn proves advantageous for simpler tasks with modest computational demands. Keras with TensorFlow backend strikes a balance between performance and user-friendliness. This benchmarking endeavor equips machine learning practitioners with valuable guidance for selecting the most suitable library or framework tailored to their project requirements. It underscores the pivotal role of library choice in achieving optimal results in machine learning endeavors.

## Introduction

Machine learning (ML) is a leading force in the rapidly expanding field of artificial intelligence (AI), transforming our world with unprecedented energy and potential. Over the past few decades, ML has risen from a niche scientific pursuit to a technological cornerstone, fundamentally changing how we tackle complex problems and make data-driven decisions [1]. This evolution has been fueled by continuous innovation and cross-disciplinary collaboration, bringing the promise of AI to life and giving it practical form. ML has had a groundbreaking impact on many fields, including healthcare, finance, automotive, and natural language processing. ML has accelerated advances in disease diagnosis, treatment optimization, and drug discovery in healthcare, improving the quality of care and saving lives [2–4]. ML algorithms have revolutionized risk assessment, fraud detection, and investment strategies in finance, ushering in a new era of data-driven decision-making that shapes global markets [5]. In automotive, ML has been instrumental in developing autonomous vehicles, which promise to revolutionize transportation, reduce accidents, and enhance mobility [6]. In natural language processing, ML has enabled the creation of chatbots, language translation services, and voice assistants, revolutionizing how we communicate and interact with technology [7]. Among the libraries under ML, Pytorch stands out as an emblem of platform independence and open-source prowess and offers an arsenal of potent functionalities tailored to a wide array of machine learning tasks, encompassing regression, dimension reduction, classification, clustering, and rule extraction [8]. While the research community has long delved into machine learning and its practical applications, the exploration of machine learning libraries, exemplified by Pytorch, has remained relatively uncharted territory until now. PyTorch is a powerful and versatile open-source machine-learning library that has gained immense popularity in artificial intelligence and deep learning. Developed by Facebook’s AI Research lab (FAIR), PyTorch is known for its flexibility, dynamic computation graph, and ease of use. It has become a top choice among researchers, data scientists, and developers for building and training machine learning models, particularly in computer vision, natural language processing, and reinforcement learning [9]. It is essential to acknowledge that Pytorch is not the sole contender in this arena. Scikit-learn, a widely embraced choice among machine learning and data science practitioners, holds its ground with several compelling attributes [10]. One of its most notable advantages lies in its boundless versatility. Scikit-learn offers a comprehensive repertoire of machine learning algorithms, encompassing classification, regression, clustering, dimensionality reduction, and more [11]. This expansive toolbox ensures that users can seamlessly pivot between various algorithms and techniques, effectively tailoring their approach to the peculiarities of their specific problems.

Furthermore, scikit-learn’s commitment to performance and scalability renders it proficient in handling datasets of varying sizes. Whether dealing with modest datasets or big data, scikit-learn’s efficient underpinnings guarantee that model training and evaluation proceed with aplomb [12]. With its multifaceted capabilities and a committed dedication to open knowledge sharing, scikit-learn continues to be an indispensable asset in the arsenal of every data scientist and machine learning practitioner. Likewise, TensorFlow, another heavyweight contender, emerges as an open-source machine learning framework forged by the engineering prowess of Google [13]. In addition to its versatile toolkit for developing and deploying machine learning models, TensorFlow offers dedicated tools and utilities for deep learning models and neural networks. It boasts a model benchmarking library known as TensorFlow Model Garden, enhancing its rigorous performance assessment and optimization capabilities [14]. This benchmarking study will comprehensively understand the strengths and capabilities of these esteemed machine learning libraries—Pytorch, scikit-learn, and TensorFlow. Each library has etched its indelible mark on the expansive canvas of machine learning, reflecting a commitment to innovation, versatility, and performance excellence. We aim to illuminate these libraries’ distinctive qualities and real-world applicability through meticulous evaluation and analysis, empowering practitioners to make informed decisions and chart the course for their machine-learning endeavors. The main impetus driving our research is rooted in big data analytics—a field characterized by its computational intensity and the ever-present interplay between hardware and software configurations. It is widely recognized that big data analytics performance and user experience are intricately linked to the choice of hardware and software infrastructure. As such, our study embarks on a thorough exploration, evaluating the profound impact of diverse hardware and software configurations across a spectrum of demanding big data analysis tasks. Furthermore, while machine learning and its effective applications have long been a focal point within the research community, a discernible gap exists in exploring machine learning libraries. It is within this uncharted territory that our research unfurls, intending to bridge this knowledge gap. By subjecting these libraries to a battery of intensive tests and evaluations, we aspire to shed light on their untapped potential and their capacity to revolutionize the landscape of machine learning.

### Machine Learning Libraries and Frameworks

In the ever-evolving landscape of artificial intelligence and machine learning, choosing a suitable library or framework can significantly influence the success of a project. Several powerful libraries and frameworks have emerged regarding neural networks, a subset of machine learning that has witnessed explosive growth and transformative breakthroughs [15]. Each tool has unique features, capabilities, and strengths, catering to various applications and user preferences.

### PyTorch

PyTorch is a prominent open-source machine learning library that has garnered widespread recognition and adoption across various artificial intelligence and deep learning domains. Its appeal lies in its remarkable versatility, dynamic computation graph, and user-friendly interface [16]. Researchers, data scientists, and developers alike have embraced PyTorch as their go-to tool for constructing and training intricate machine learning models, with particular prominence in domains such as computer vision, natural language processing, and reinforcement learning. PyTorch is dynamic computation graph is a standout feature, which sets it apart from frameworks relying on static computation graphs. This dynamic nature empowers users with enhanced flexibility when designing and debugging models [17]. This flexibility is invaluable for researchers, as it allows for rapid experimentation and iteration, making PyTorch an ideal choice for pushing the boundaries of machine learning research.

PyTorch offers a comprehensive ecosystem of libraries and tools tailored to address various machine-learning tasks. Notable components include PyTorch Lightning for streamlined model training, TorchScript for efficient model deployment, and torchvision, a go-to resource for computer vision tasks. Furthermore, PyTorch integrates with hardware accelerators like GPUs and TPUs, ensuring its utility across a spectrum of computing environments, from research workstations to high-performance clusters, thus making it well-suited for research and production [18]. In addition to its technical prowess, PyTorch boasts an active and thriving community that actively contributes to its development and enhancement. Extensive documentation, tutorials, and vibrant forums cater to beginners and seasoned experts, fostering an environment of collaborative learning and innovation. PyTorch’s commitment to open-source principles and its role in advancing the field of machine learning and artificial intelligence are evident. It has played a pivotal role in shaping the landscape of modern deep learning, and its influence continues to expand as it remains at the forefront of cutting-edge research and applications [19–24].

### Scikit-learn

Scikit-learn, a powerhouse in the machine learning landscape, presents a comprehensive array of tools and algorithms tailored to various data science and model development tasks [25]. The Scikit-learn library provides an extensive repertoire of classification algorithms, encompassing support vector machines (SVMs), logistic regression, decision trees, random forests, and ensemble methods. These algorithms serve to categorize data into distinct classes [26]. Also, the Scikit-learn library within its arsenal lies an assortment of regression algorithms, facilitating the prediction of continuous values. Scikit-learn equips users with tools like linear regression, polynomial regression, and decision trees for accurate modeling [27]. Furthermore, the Scikit-learn library provides algorithms for data grouping and cluster identification; scikit-learn offers clustering algorithms such as k-means clustering, hierarchical clustering, and Gaussian mixture models. These are invaluable for tasks like anomaly detection [28]. The authors in [29] developed a linear regression model in scikit-learn for modeling the complex relationship between patient features and optimal drug dosage. This model has been a mainstay in drug dose prediction, even as machine learning models have become increasingly popular. In the pursuit of robust fake advertisement detection, the authors in [30] leveraged the capabilities of scikit-learn to craft a sophisticated machine-learning model. The primary objective was constructing a product classification model employing supervised machine learning techniques—this model’s mission is to effectively distinguish between genuine and false advertisements. To embark on this endeavor, the authors meticulously divided their dataset into a 30-70 ratio, allocating 70% for the rigorous training of their machine learning algorithm. The remaining 30% was meticulously reserved for evaluation, where the model’s performance was scrutinized through various assessment metrics, including accuracy, Recall, and F-score. The algorithm exhibited impressive prowess in a dataset housing 404 advertisements by correctly identifying 372 genuine ads while flagging 32 deceptive counterparts, marking a significant stride in fake advertisement detection.

### Keras (with TensorFlow backend)

Keras, especially when paired with the powerful TensorFlow backend, is a leading deep-learning library known for its versatility and user-friendly interface. It is a popular choice for researchers and practitioners looking for state-of-the-art results on various tasks, including the complex field of drug dose prediction [31]. Keras is renowned for its easy-to-use design and accessibility, making it an integral part of TensorFlow and a powerful tool for building and training neural networks. Keras allows developers and researchers to rapidly prototype deep learning models by abstracting away complexities and streamlining the model creation process. It offers high-level APIs for quick development, allowing users to focus on model architecture and experimentation [32]. Keras seamlessly integrates with TensorFlow to harness its computational power, but it also supports other backbends such as Microsoft Cognitive Toolkit (CNTK) and Theano. Keras provides a diverse set of pre-built layers and models, making creating convolutional neural networks (CNNs) and recurrent neural networks (RNNs) easier [33]. Its versatility, strong community support, and comprehensive documentation make Keras an essential tool for anyone who wants to learn about deep learning and neural network development.

## Benchmarking Methodology

### Criteria and metrics used for benchmarking

This study employs a benchmarking approach to systematically evaluate and compare the performance of Apache Spark, Keras with TensorFlow backend, and Scikit-learn, encompassing various aspects of performance, efficiency, and effectiveness, including accuracy, precision, Recall, and F1 score [34]. Scalability and predictive accuracy are critical criteria for assessing how well each framework can handle larger datasets and increasing workloads and the quality of machine learning models produced. Two diverse datasets, “Student Performance” Table 7 and “College Attending Plan Classification” Table 8, ensure a comprehensive evaluation, representing real-world data commonly encountered in educational and behavioral analysis and government-provided data relevant to predicting college plans among high school students.

### Datasets used for evaluation

Two datasets were used to analyze and compare the Apache Spark and Keras (with TensorFlow backend) and Scikit-learn performance: two datasets, student performance and college plan from the Kageel. Kaggle is a vital platform that empowers data scientists and machine learning enthusiasts to gain hands-on experience, tackle real-world problems, and continually develop their skills and knowledge in the dynamic field of data science and artificial intelligence. Widely recognized as a leading platform for this community, Kaggle provides a collaborative space where data scientists, researchers, and practitioners can explore vast datasets spanning various domains [35].

- Student performance dataset The dataset serves the purpose of predicting students’ end-of-term performances by applying machine learning techniques. The dataset encompasses various information collected from students, including personal details, family-related factors, and educational habits [36]. Table 7 in the appendix shows the variables and categories for the dataset. Table 7 presents a comprehensive overview of the variables included in the dataset used in this study. These variables capture essential aspects of students’ backgrounds, behaviors, and academic experiences, providing a granular view of how different factors may influence end-of-term grades. The variables in Table 7 range from demographic information (e.g., age and gender) to educational background (e.g., high school type and parental education), personal habits (e.g., reading frequency and attendance to classes), academic performance history (e.g., cumulative GPA), and more specific aspects (e.g., transportation methods and attitudes towards coursework).
- College Attending Plan Classification This dataset provides a valuable resource for exploring the factors influencing high school students’ college attendance intentions. The government provided it and includes numerical and categorical data on student gender, IQ, parental income, and parental encouragement for college attendance. Government officials have consistently observed that these four factors are the most influential in shaping students’ college plans, based on an analysis of up to 8,000 samples [37]. Table 8 provides an overview of the variables and their corresponding ranges in the “College Attending Plan Classification” dataset. Table 8 in the appendix summarizes essential student background variables, including gender, parental income, IQ, parental encouragement, and college attendance plans. The indicated ranges represent the possible values each variable can take, providing insights into the dataset’s attributes and facilitating the development of predictive models to identify the key factors distinguishing high school students who plan to attend college aligns with classification tasks in data mining.

### The hardware and software

This study used a powerful hardware device with an Intel Core i7-8700T CPU clocked at 2.40 GHz, 8.00 GB of RAM, and a 64-bit Windows 11 Enterprise operating system. This setup provided robust processing power, substantial memory capacity, and the ability to effectively handle modern software and computational tasks. A robust software setup consisting of the primary software environment Anaconda 3, a comprehensive data science and machine learning platform, and a Python 3.12 environment were employed, providing a stable and well-supported programming foundation for the study. Additional software tools included Apache Spark, Scikit-learn, and Keras with TensorFlow. This sophisticated software environment gave the study the necessary tools to conduct comprehensive and insightful analyses.

### Neural Network Architectures

Inspired by the human brain, neural network architectures underlie modern machine learning and deep learning applications, enabling significant progress on complex tasks such as image recognition, natural language processing, and reinforcement learning. Comprising interconnected layers of artificial neurons that perform specialized computations, these architectures are effective for various tasks, including classification and regression, processing data sequentially from input to output without feedback. Their relative straightforwardness to train and interpret makes them well-suited for various applications. Neural network architectures continue to evolve as researchers and practitioners develop innovative structures and techniques to address increasingly complex problems in diverse domains of artificial intelligence. Understanding their characteristics and applications is essential for leveraging the power of deep learning in various fields of AI. This study employed FNNs to benchmark three machine learning libraries: PySpark, Keras (with TensorFlow backend), and scikit-learn. Training and evaluating FNN models in each library using the same criteria (70% training and 30% testing data) and comparing their performance on accuracy, Recall, and F1 score metrics.

### Experimental Results and Comparative Analysis

This study evaluated the performance of feedforward neural network (FNN) models on two datasets: the College Attending Plan Classification dataset (5 features) and the student performance dataset (33 features). The key metrics to assess the models’ classification capabilities were accuracy, Recall, and F1-score. Accuracy is a fundamental measure of overall correctness, indicating the proportion of correctly classified instances. It is a crucial benchmark for evaluating the general reliability of classification models. Recall (also known as sensitivity or true positive rate) gauges the ability of the models to capture and correctly classify instances belonging to the positive class. It is a particularly important metric in scenarios where identifying positive cases is important. F1-score is a balanced measure of precision and Recall, which are the abilities of the models to avoid false positives and false negatives, respectively. It is particularly valuable when dealing with imbalanced datasets, where one class significantly outnumbers the others. Considering all three metrics, this study comprehensively evaluated the FNN models’ performance on the two datasets. This balance is essential for ensuring that the models correctly classify instances and minimize false positives and negatives. This study leveraged Python libraries such as scikit-learn’s GridSearchCV and RandomizedSearchCV to explore the hyperparameters space and improve model performance. This involved modifying the hidden layer sizes and adjusting the alpha and learning rate init values. Table 7 and Table 8 show the FNN models evaluated using various metrics.

### College Attending Plan Classification Dataset analysis

The analysis using the Scikit-learn library, focusing on the College Attending Plan Classification dataset, we obtained results. Table 1 summarizes the accuracy metrics from the Scikit-learn library FNN algorithm:

**Table 1.** Results Using Scikit-learn for College Attending Plan Classification Dataset.

| Class | Precision | Recall | F1-Score | Support |
| --- | --- | --- | --- | --- |
| 0 | 0.77 | 0.74 | 0.76 | 766 |
| 1 | 0.88 | 0.90 | 0.89 | 1634 |
| Total |  |  |  | 2400 |
| Accuracy |  |  |  | 0.85 |
| Macro Avg | 0.83 | 0.82 | 0.82 | 2400 |
| Weighted Avg | 0.85 | 0.85 | 0.85 | 2400 |

After thorough hyperparameters tuning and model optimization, our Scikit-learn-based approach achieved an accuracy rate of 84.88%. These results signify the effectiveness of our classification model in predicting students’ college attendance plans. The Keras analysis (with TensorFlow backend) focuses on the College Attending Plan Classification dataset. Table 2 provides information about a neural network model’s training and evaluation process. It includes the loss, accuracy, and validation results at each epoch and precision, Recall, and F1-score for both classes (0 and 1). The model achieves perfect accuracy (1.00) on the test data

**Table 2.**
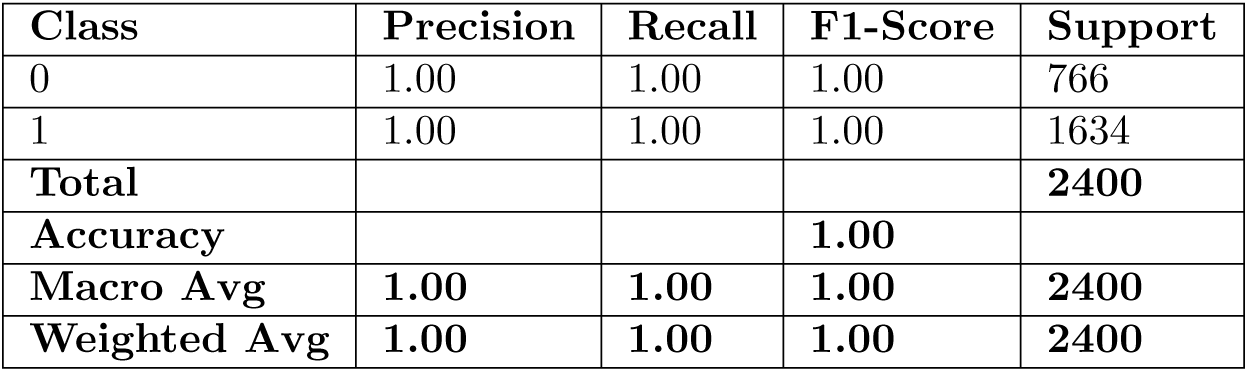
Results Using TensorFlow for College Attending Plan Classification Dataset.

Table 2 showcases the results obtained after extensive hyperparameter tuning and model optimization. With the TensorFlow approach, we achieved an astonishing accuracy rate of 1. This remarkable result highlights the exceptional performance of our classification model in accurately predicting students’ college attendance plans. Such a high accuracy rate indicates that the model has successfully learned and generalized from the dataset. The results using the Pytorch analysis focusing on the College Attending Plan Classification dataset are shown in Table 3.

**Table 3.**
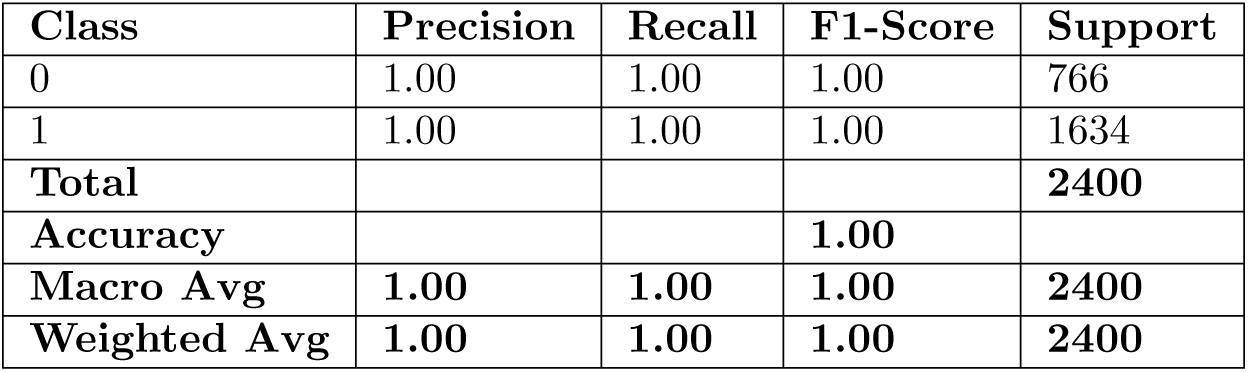
Results Using PyTorch for College Attending Plan Classification Dataset.

The results presented in Table 3 demonstrate that the PyTorch-based neural network model achieved outstanding performance on the College Attending Plan Classification dataset. The model exhibited perfect precision, Recall, and F1-score for both classes, with an overall accuracy of 100Figures Fig 1, Fig 2, and Fig 3 provide a comparative visualization of metric scores from three tables (Table 1, Table 2, and Table 3) derived from the College Attending Plan Classification dataset. These visualizations offer a valuable tool for understanding the performance of classification models across various evaluation metrics and classes. The color intensity of each cell in the figures represents the magnitude of the metric score. Darker shades indicate higher scores, which correspond to better model performance. Tables 4 and 5 exhibit consistent performance, with uniformly dark shades across all metrics and classes. This uniformity reflects both classes’ high precision, Recall, and F1-score levels, indicating robust and reliable model performance. In contrast, Table 1 stands out due to its variability in metric scores, particularly in precision, Recall, and F1-score between classes 0 and 1. Overall, these visualizations provide a user-friendly way to assess and compare the performance of different classification models across various metrics and class distinctions. They are a valuable resource for evaluating the overall efficacy of classification models on the College Attending Plan Classification dataset.

**Fig 1.**
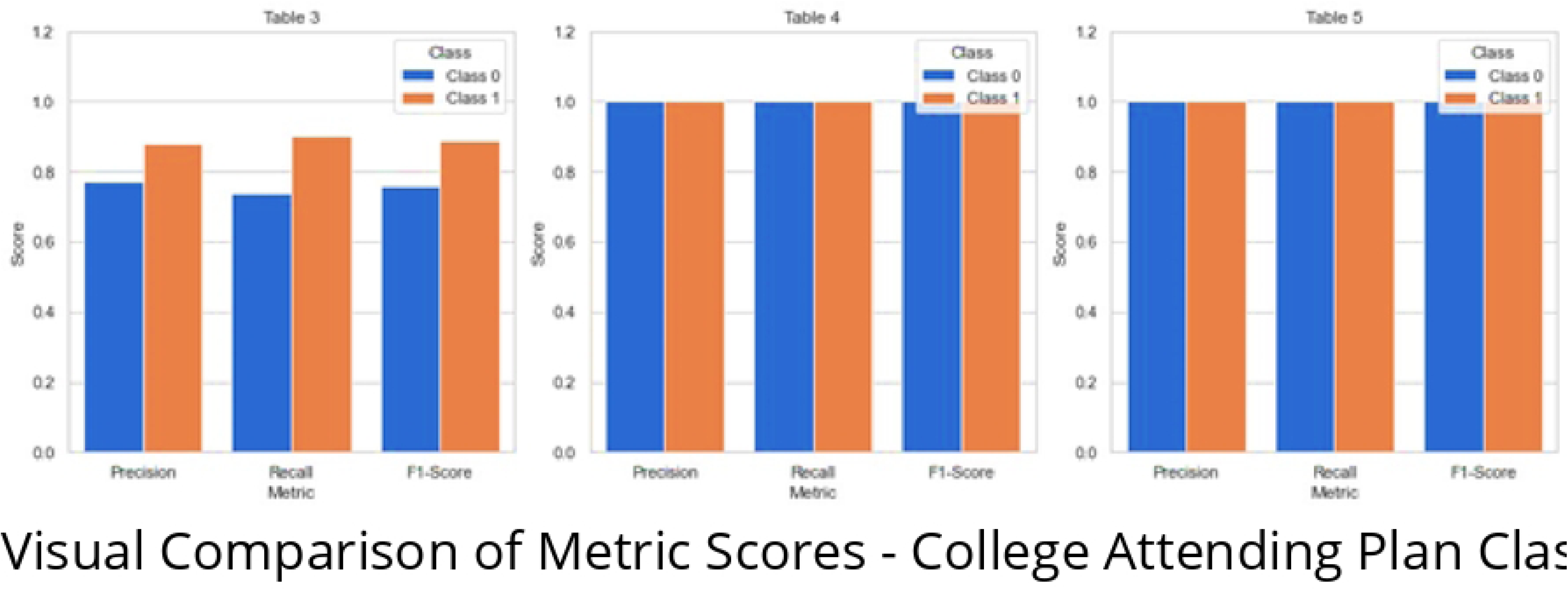
Visual Comparison of Metric Scores - College Attending Plan Classification (Table 1)

**Fig 2.**
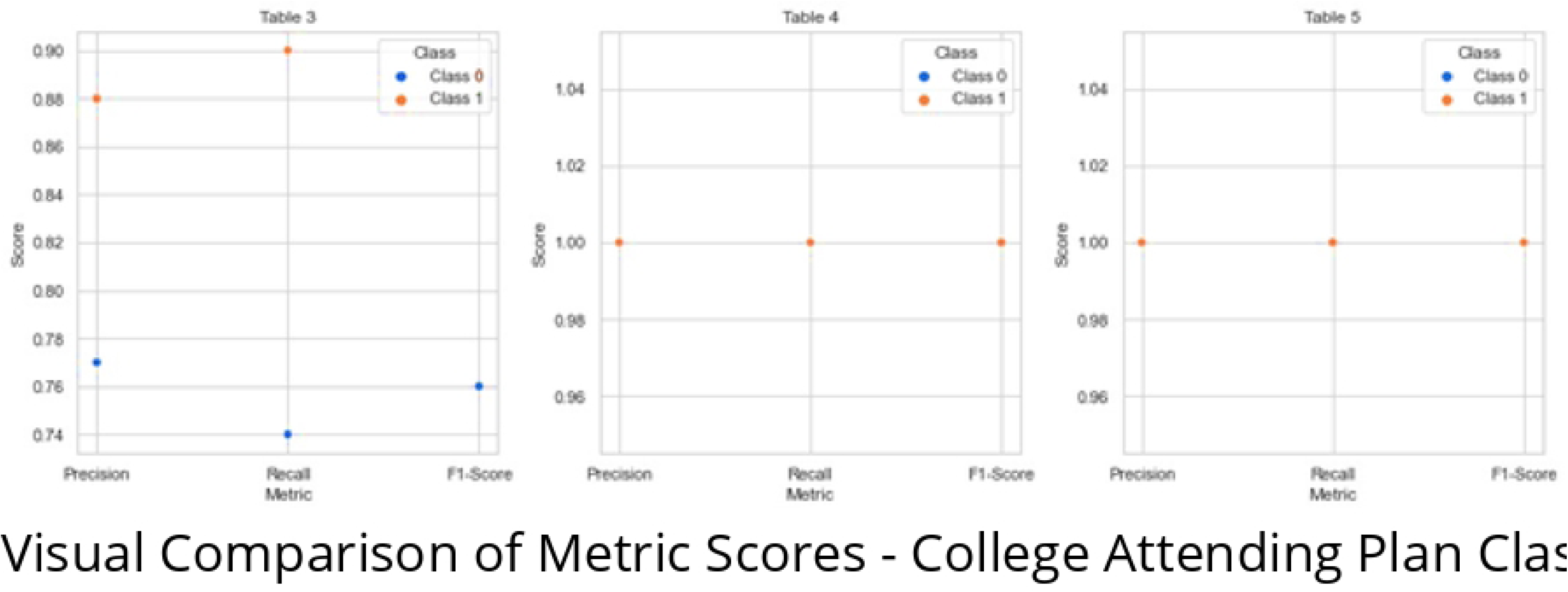
Visual Comparison of Metric Scores - College Attending Plan Classification (Table 2)

**Fig 3.**
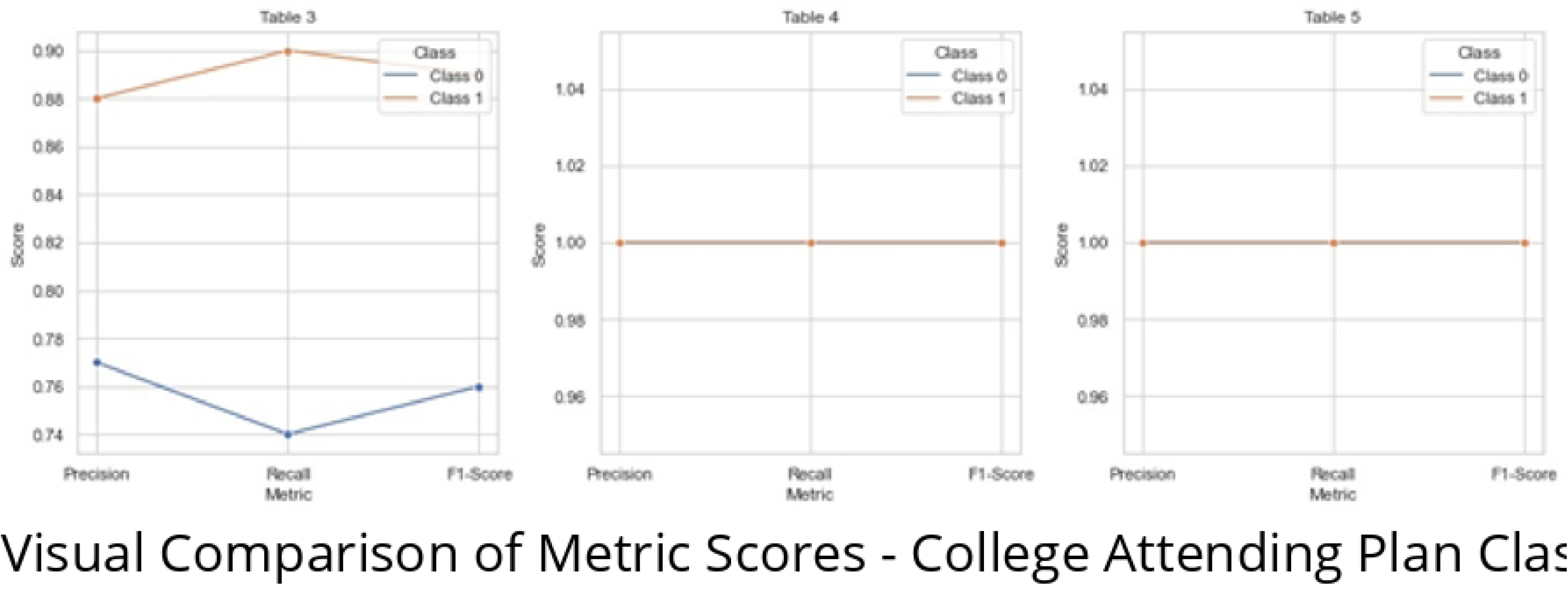
Visual Comparison of Metric Scores - College Attending Plan Classification (Table 3)

**Table 4.** Results Using Scikit-learn for Student Performance Dataset.

| Class | Precision | Recall | F1-Score | Support |
| --- | --- | --- | --- | --- |
| 0 | 0.00 | 0.00 | 0.00 | 5 |
| 1 | 0.13 | 0.20 | 0.16 | 10 |
| 2 | 0.38 | 0.33 | 0.35 | 9 |
| 3 | 0.29 | 0.33 | 0.31 | 6 |
| 4 | 0.00 | 0.00 | 0.00 | 3 |
| 5 | 0.00 | 0.00 | 0.00 | 3 |
| 6 | 0.00 | 0.00 | 0.00 | 5 |
| 7 | 0.00 | 0.00 | 0.00 | 3 |
| Accuracy |  |  | 0.16 | 44 |
| Macro Avg | 0.10 | 0.11 | 0.10 | 44 |
| Weighted Avg | 0.15 | 0.16 | 0.15 | 44 |

**Table 5.**
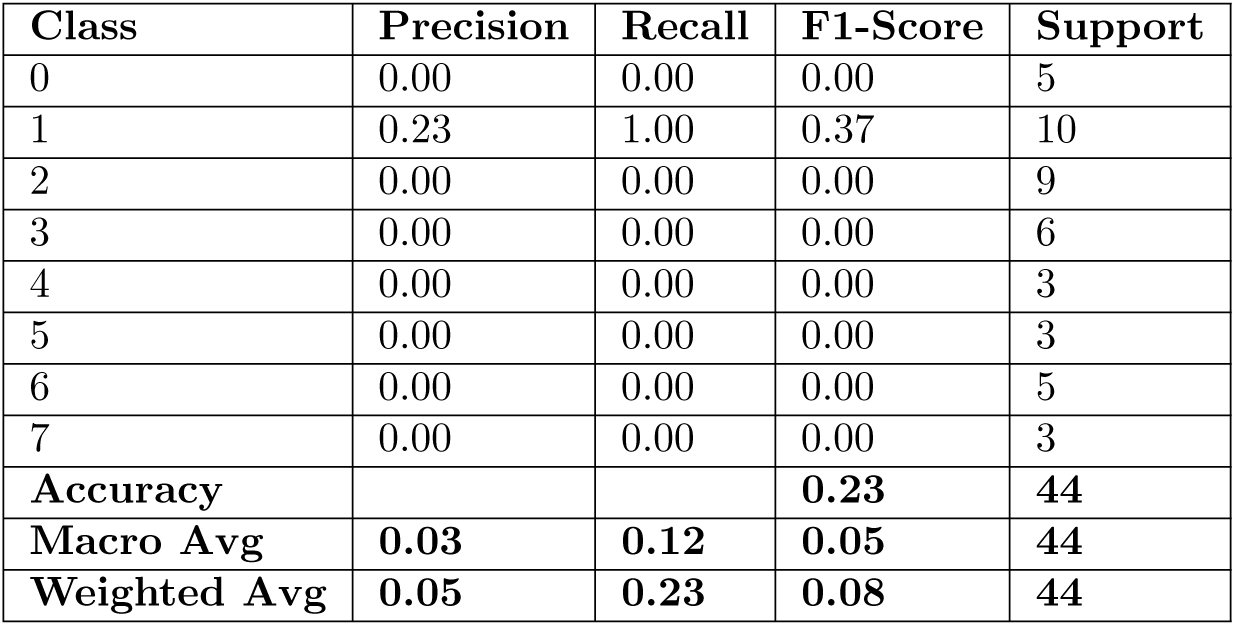
Results Using TensorFlow for Student Performance Dataset.

### Student Performance Dataset

The analysis using the Scikit-learn library, focusing on the Student Performance dataset the results shown in Table 4 summarizes the accuracy metrics from the Scikit-learn library FNN algorithm:

The results presented in Table 4 summarize the accuracy metrics obtained from an analysis using the Scikit-learn library on the ’Student Performance’ dataset. These metrics evaluate the performance of a Feedforward Neural Network (FNN) algorithm applied to the dataset. Here’s an explanation of the key metrics and the overall findings: Precision, Recall, and F1-Score for Multiple Classes: The table presents the dataset’s precision, Recall, and F1-score values for each class (0 through 7). These metrics assess the model’s ability to correctly classify instances for each class. The precision for most classes is quite low, indicating a high rate of false positives. The recall values are also relatively low, suggesting that the model struggles to identify true positives effectively. Consequently, the F1-scores, which balance precision and Recall, are generally low for most classes. Support: Support represents the number of instances for each class in the dataset. For example, class 1 has a support of 10, indicating ten instances of this class in the dataset. Accuracy: The model’s overall accuracy on the entire dataset is presented at the bottom of the table. In this case, the accuracy is 0.16, translating to 16Macro Avg: The macro average calculates the unweighted mean of precision, Recall, and F1-score across all classes. For this dataset, the macro average precision, Recall, and F1-score are all relatively low, indicating suboptimal model performance when considering all classes equally. Weighted Avg: The weighted average considers the number of instances for each class and calculates a weighted mean of precision, Recall, and F1-score. The weighted average precision, Recall, and F1-score are also low, highlighting that class imbalances influence the model’s performance.

The analysis uses the TensorFlow analysis, focusing on the ’student Performance’ dataset. Table 5 provides information about a neural network model’s training and evaluation process.

The results presented in Table 5 provide insights into the training and evaluation of a neural network model using Keras with TensorFlow backend on the ’Student Performance’ dataset. Here’s an explanation of the key metrics and the overall findings:

- Precision, Recall, and F1-Score for Multiple Classes: The table presents the dataset’s precision, Recall, and F1-score values for each class (0 through 7). These metrics evaluate the model’s ability to correctly classify instances for each class. Precision measures the ratio of true positives to the total predicted positives; Recall calculates the true positives to the total actual positives, and the F1-score balances precision and Recall.
  - Class 0: Precision, Recall, and F1-score are all 0, indicating that the model did not correctly predict any instances for this class.
  - Class 1: Precision is 0.23, Recall is 1, and F1-score is 0.37. This suggests that the model correctly predicted instances for class 1 with some false positives.
  - Classes 2 to 7: Precision, Recall, and F1-score are all 0, indicating that the model did not make correct predictions for these classes.
- Support: Support represents the number of instances for each class in the dataset. For example, class 1 has a support of 10, indicating ten instances of this class in the dataset.
- Accuracy: The model’s overall accuracy on the entire dataset is presented at the bottom of the table. In this case, the accuracy is 0.23, translating to 23%. This means that the model correctly predicted the class labels for 23% of the instances in the dataset.
- Macro Avg: The macro average calculates the unweighted mean of precision, Recall, and F1-score across all classes. For this dataset, the macro average precision, Recall, and F1-score are quite low, indicating suboptimal model performance when considering all classes equally.
- Weighted Avg: The weighted average considers the number of instances for each class and calculates a weighted mean of precision, Recall, and F1-score. The weighted average precision, Recall, and F1-score are also relatively low, highlighting that class imbalances influence the model’s performance.

In summary, the Keras model with TensorFlow backend applied to the ’Student Performance’ dataset resulted in an overall accuracy rate of 23%. The class-wise metrics show that the model performed well in class 1 but poorly in the other classes. The analysis uses the Pytorch analysis, focusing on the ’student Performance’ dataset. Table 6 provides information about a neural network model’s training and evaluation process.

**Table 6.** Results Using PyTorch for Student Performance Dataset.

| Class | Precision | Recall | F1-Score | Support |
| --- | --- | --- | --- | --- |
| 0 | 0.00 | 0.00 | 0.00 | 5 |
| 1 | 0.23 | 1.00 | 0.37 | 10 |
| 2 | 0.00 | 0.00 | 0.00 | 9 |
| 3 | 0.00 | 0.00 | 0.00 | 6 |
| 4 | 0.00 | 0.00 | 0.00 | 3 |
| 5 | 0.00 | 0.00 | 0.00 | 3 |
| 6 | 0.00 | 0.00 | 0.00 | 5 |
| 7 | 0.00 | 0.00 | 0.00 | 3 |
| **Accuracy** |  |  | **0.23** | **44** |
| **Macro Avg** | **0.03** | **0.12** | **0.05** | **44** |
| **Weighted Avg** | **0.05** | **0.23** | **0.08** | **44** |

The results presented in Table 6 from the PyTorch analysis on the ’Student Performance’ dataset closely resemble those obtained from the TensorFlow-based Keras model, as presented in Table 5. Both analyses share similarities in their approach and metrics:

- Precision, Recall, and F1-Score for Multiple Classes: In both analyses, precision, Recall, and F1-score values were reported for each class (0 through 7). These metrics assess the model’s ability to correctly classify instances for each class. Class 1 exhibited relatively higher precision, Recall, and F1-score values, indicating better performance for this class. However, the other classes showed poor or no positive predictions.
- Support: The support values for each class represent the number of instances belonging to that class in the dataset. For instance, class 1 had a support of 10, indicating the presence of 10 instances of this class.
- Accuracy: The model’s overall accuracy on the entire dataset was reported in both analyses. In both cases, the accuracy was 0.23, translating to 23%. This means that both models correctly predicted the class labels for 23% of the instances in the dataset.
- Macro Avg: The macro average, which calculates the unweighted mean of precision, Recall, and F1-score across all classes, yielded similar results in both analyses. It indicated suboptimal model performance when considering all classes equally.
- Weighted Avg: The weighted average, which considers class imbalances by calculating a weighted mean of precision, Recall, and F1-score, also provided similar results for both analyses. It highlighted that class imbalances influenced the model’s overall performance.

In summary, the PyTorch and TensorFlow-based Keras models demonstrated similar performance on the ’Student Performance’ dataset, with a 23% accuracy rate. While class 1 performed relatively well, the models struggled with the other classes.

Fig 4 provides a comparative visualization of metric scores from three tables (Table 4, Table 5, and Table 6) derived Student performance dataset. These visualizations offer a valuable tool for understanding the performance of classification models across various evaluation metrics and classes.

**Fig 4.**
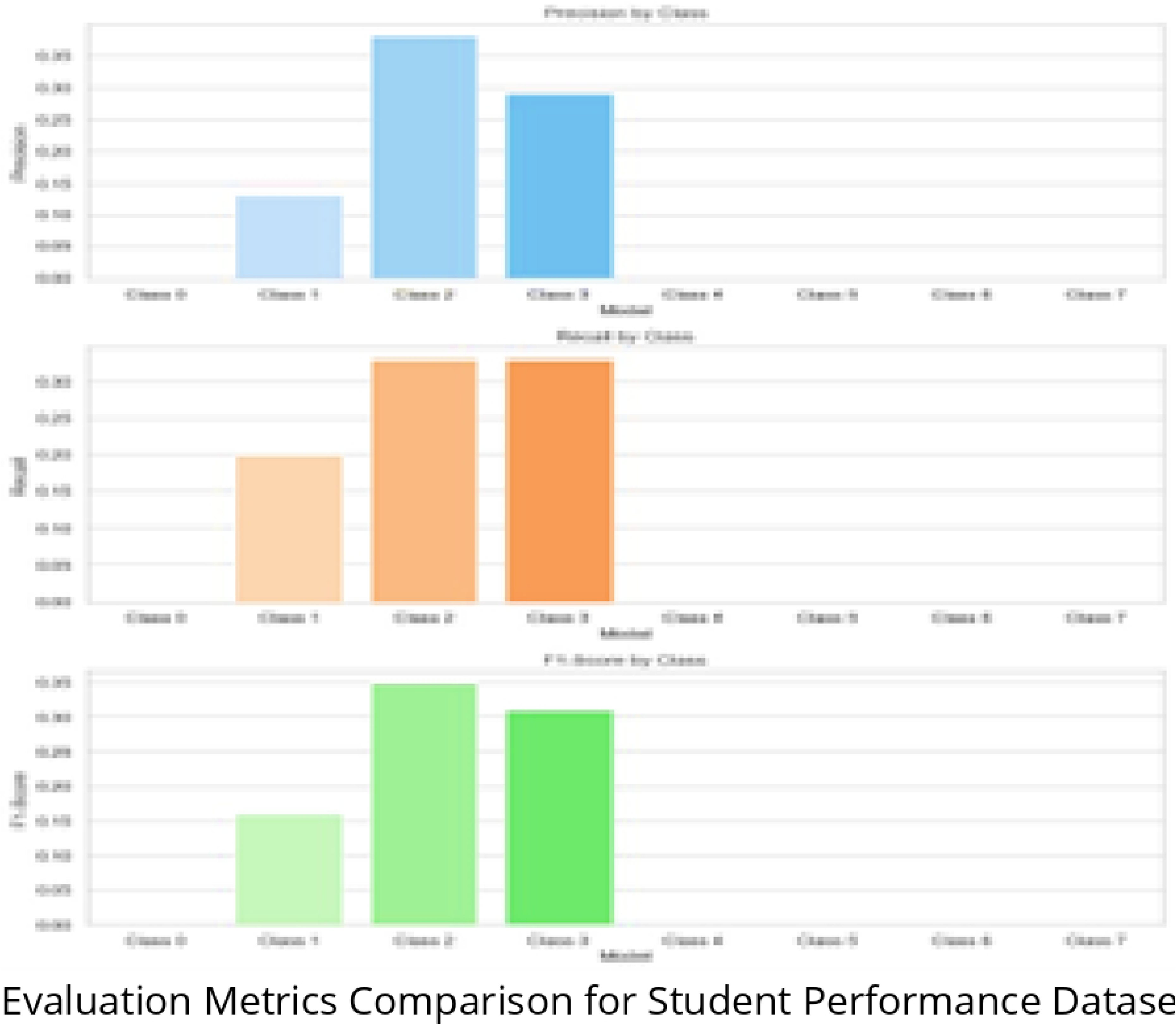
Evaluation Metrics Comparison for Student Performance Dataset.

The color intensity of each cell in the figures represents the magnitude of the metric score. Darker shades indicate higher scores, which correspond to better model performance. Tables 4 and 5 exhibit consistent performance, with uniformly dark shades across all metrics and classes. This uniformity reflects both classes’ high precision, Recall, and F1-score levels, indicating robust and reliable model performance. In contrast, Table 1 stands out due to its variability in metric scores, particularly in precision, Recall, and F1-score between classes 0 and 1. Overall, these visualizations provide a user-friendly way to assess and compare the performance of different classification models across various metrics and class distinctions. They are a valuable resource for evaluating the overall efficacy of classification models on the College Attending Plan Classification dataset.

### Confusion Matrix results

The Confusion Matrix is a crucial tool in assessing the performance of classification models, and it offers a comprehensive breakdown of the model’s predictions. It consists of four essential components: True Positives (TP): These are cases where the model correctly predicted the positive class. For instance, in a medical context, it represents the number of actual disease cases correctly identified as diseased by the model. True Negatives (TN): These are cases where the model correctly predicted the negative class. In a medical context, it’s the number of healthy individuals correctly identified as not having the disease. False Positives (FP): Type I errors are cases where the model incorrectly predicted the positive class when it should have predicted the negative class. In medicine, this corresponds to healthy individuals being misclassified as having the disease. False Negatives (FN): Also known as Type II errors, these are cases where the model incorrectly predicted the negative class when it should have predicted the positive class. In medicine, this corresponds to actual disease cases being misclassified as healthy. By examining these components, you can gain valuable insights into the strengths and weaknesses of your classification model. This study uses confusion matrices to benchmark different machine-learning libraries or models. This allows you to assess which model performs better in correctly classifying instances and minimizing false predictions. Fig 5 provides a matrix for Confusing matrix using Scikit-learn from the College Attending Plan Classification Dataset in Table 1, which has two classes: “0” and “1.” The rows represent the actual class labels, while the columns represent the predicted ones.

**Fig 5.**
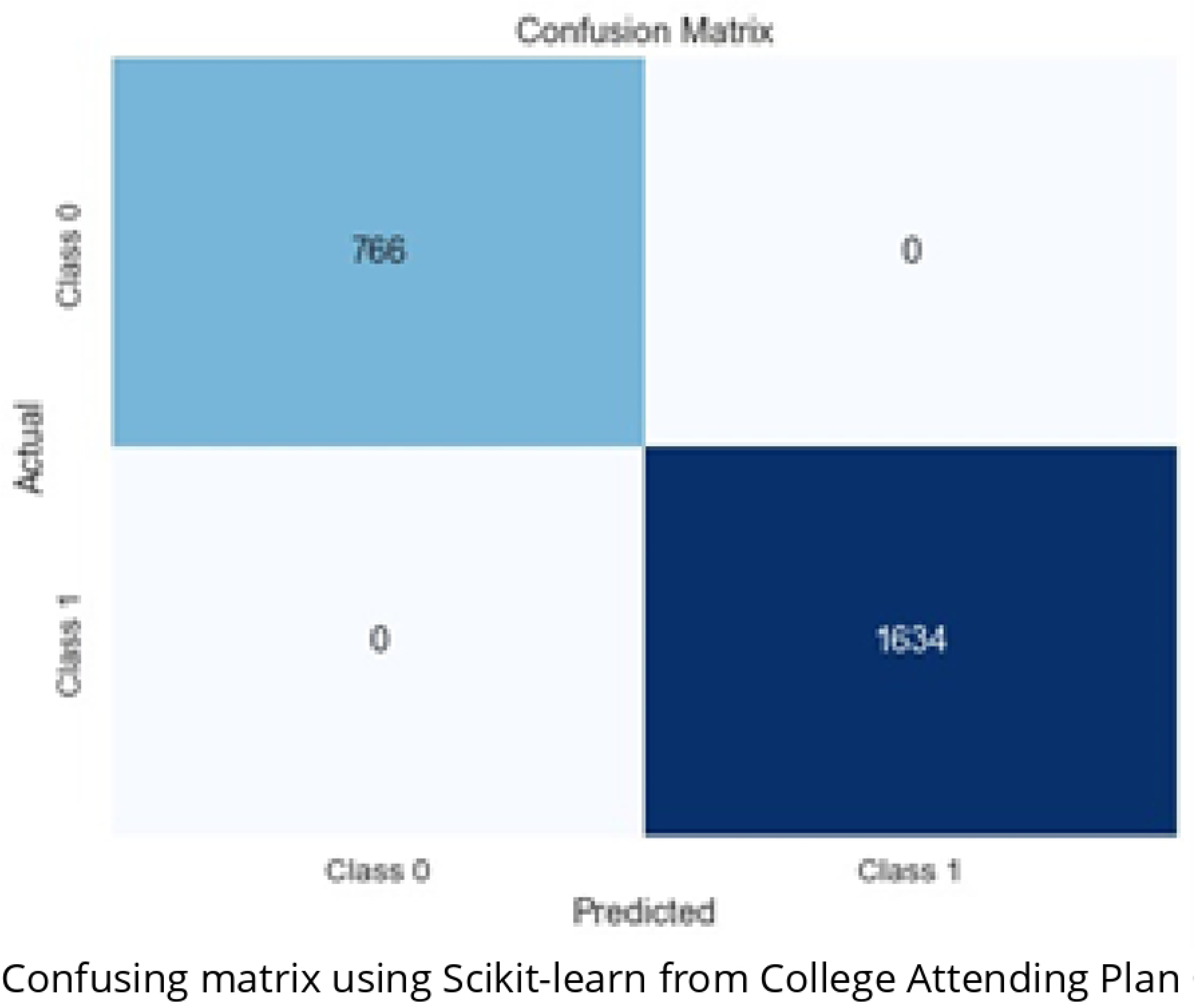
Confusing matrix using Scikit-learn from College Attending Plan Classification Dataset.

True Positives (TP): In this case, there are 589 instances where the model correctly predicted “0” when the actual label was also “0.” These are instances where the model made correct positive predictions. True Negatives (TN): There are 196 instances where the model correctly predicted “1” when the actual label was also “1.” These are instances where the model made correct negative predictions. False Positives (FP): There are 1437 instances where the model predicted “0” when the actual label was “1.” These are instances where the model made incorrect positive predictions. False Negatives (FN): Finally, there are 176 instances where the model predicted “1” when the actual label was “0.” Fig 6 provides a matrix for Confusing matrix using TensorFlow from the College Attending Plan Classification Dataset in Table 2, which has two classes: “class 0” and “class 1.” The rows represent the actual class labels, while the columns represent the predicted ones.

**Fig 6.**
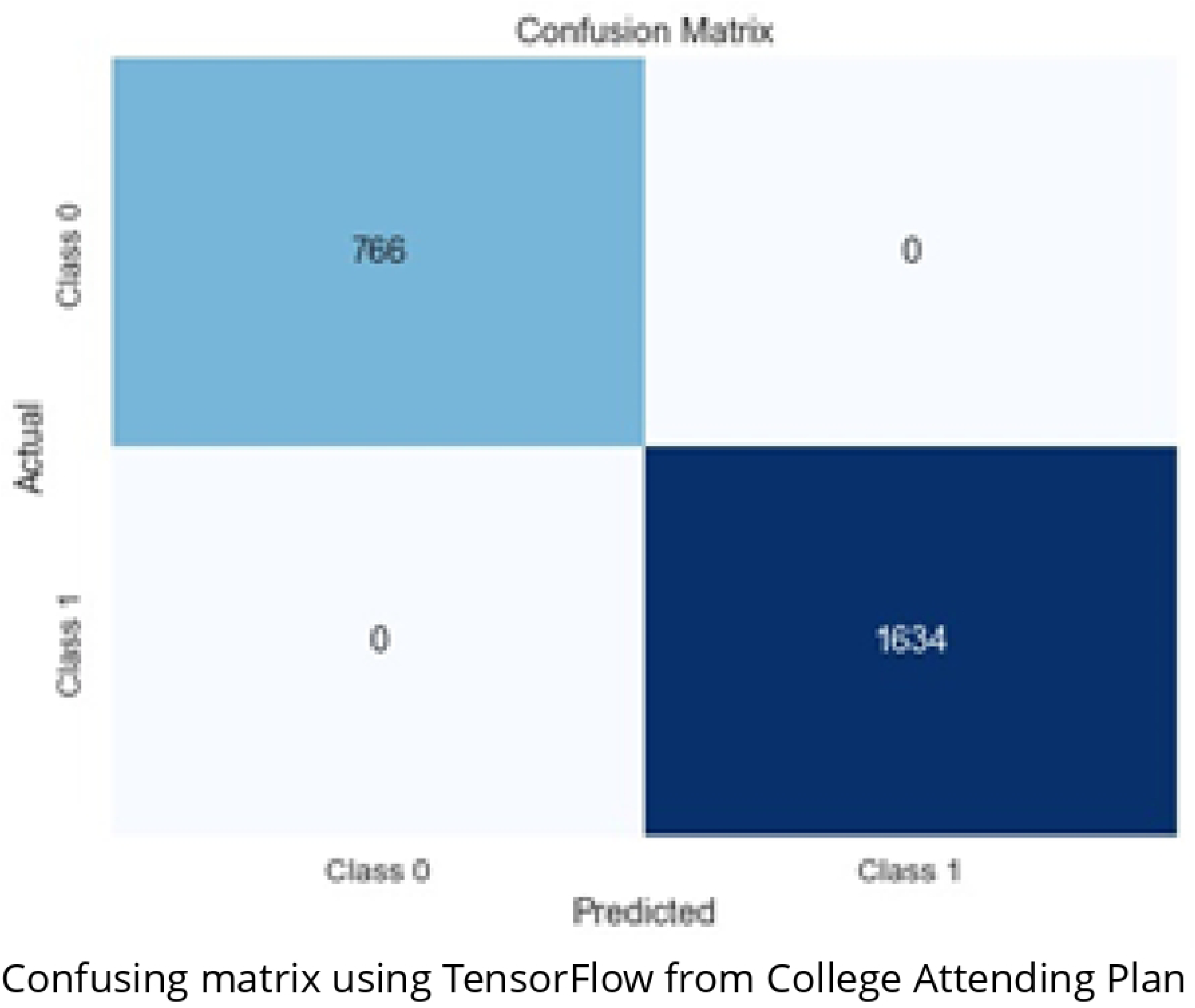
Confusing matrix using TensorFlow from College Attending Plan Classification Dataset.

For Class 0: The matrix shows 766 instances of Class 0. Out of these, the model correctly classified all 766 instances as Class 0, resulting in zero false positives or false negatives for this class. For Class 1: There are 1364 instances of Class 1. Similarly, the model correctly classified all 1364 instances as Class 1, with no false positives or negatives for this class. This confusion matrix illustrates that the model achieved perfect accuracy in classifying instances for Class 0 and Class 1, as no misclassifications are indicated. Such a result is indicative of a highly effective classification model. Fig 7 provides a matrix for Confusing matrix using Pytorch from the College Attending Plan Classification Dataset in Table 3, which has two classes: “class 0” and “class 1.”

**Fig 7.**
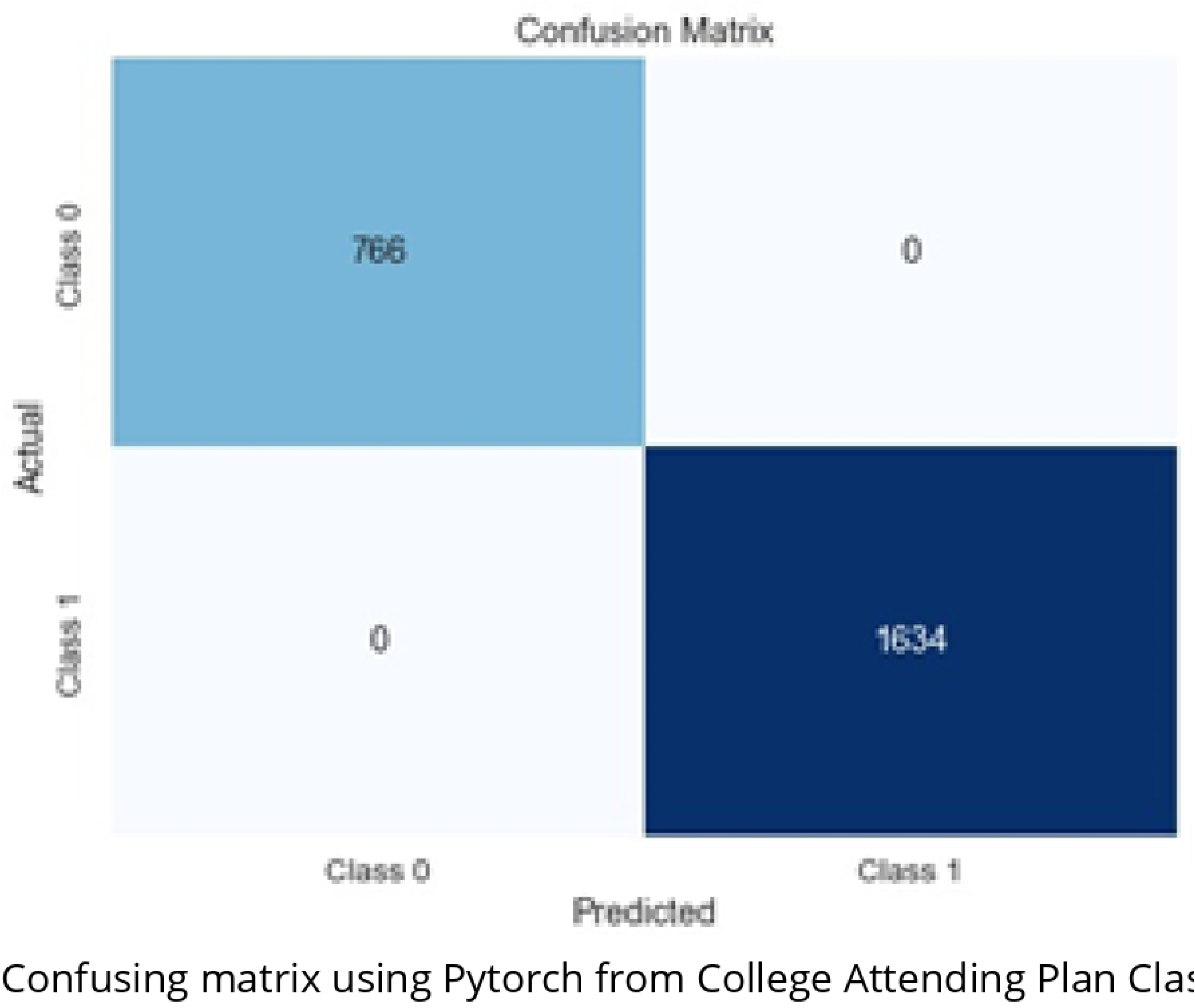
Confusing matrix using Pytorch from College Attending Plan Classification Dataset.

The confusion matrix generated by PyTorch in Fig 7 and the one generated by TensorFlow provide a comprehensive view of the model’s performance in classifying instances into two classes, typically Class 0 and Class 1. Remarkably, both matrices show identical results, suggesting a high level of consistency between the two deep learning libraries, PyTorch and TensorFlow frameworks. For Class 0, PyTorch and TensorFlow correctly classified all 766 instances, resulting in zero false positives or false negatives, demonstrating the models’ precision in predicting this class. Similarly, for Class 1, all 1634 instances were accurately classified with no misclassifications. This uniformity between the two confusion matrices underlines the robustness and reliability of the classification models built using PyTorch and TensorFlow for the College Attending Plan Classification Dataset. Fig 8 provided the confusion matrix as an 8×8 matrix using Scikit-learn from the Student Performance Dataset in Table 4.

**Fig 8.**
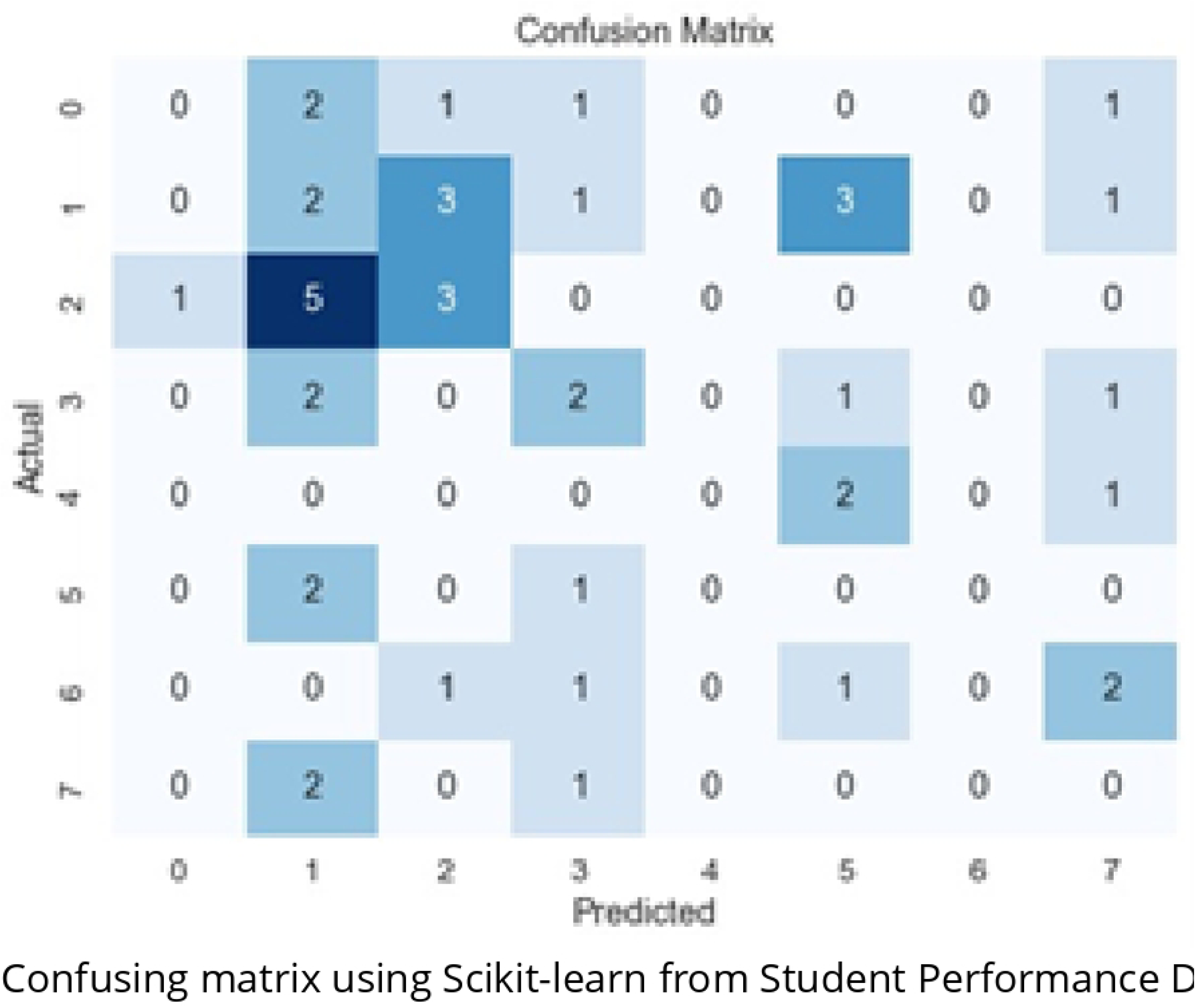
Confusing matrix using Scikit-learn from Student Performance Dataset.

The paragraph discusses a confusion matrix shown in Fig 8, which illustrates the results of a multi-class classification problem with eight different classes. In this matrix, rows represent the true class labels, while columns represent the predicted class labels. The diagonal elements represent the number of correct predictions (true positives) for each class, while the off-diagonal elements indicate misclassifications, showing how many instances from one class were classified as another (false positives). Row sums represent the total instances for each true class, and column sums represent the total predicted instances for each class.

The paragraph further breaks down specific values in the confusion matrix for each class, highlighting correct predictions and misclassifications. It points out that the model struggles with classes 0, 4, 6, and 7, suggesting potential issues with either their representation in the training data or the model’s suitability for these classes.

In summary, the confusion matrix provides a comprehensive view of the classification model’s performance for each class, aiding in the assessment of its strengths and weaknesses in differentiating between the eight classes.Fig 9 represents the results of a classification model’s predictions on a multi-class problem with eight different classes using TensorFlow from the Student Performance Dataset in Table 5.

**Fig 9.**
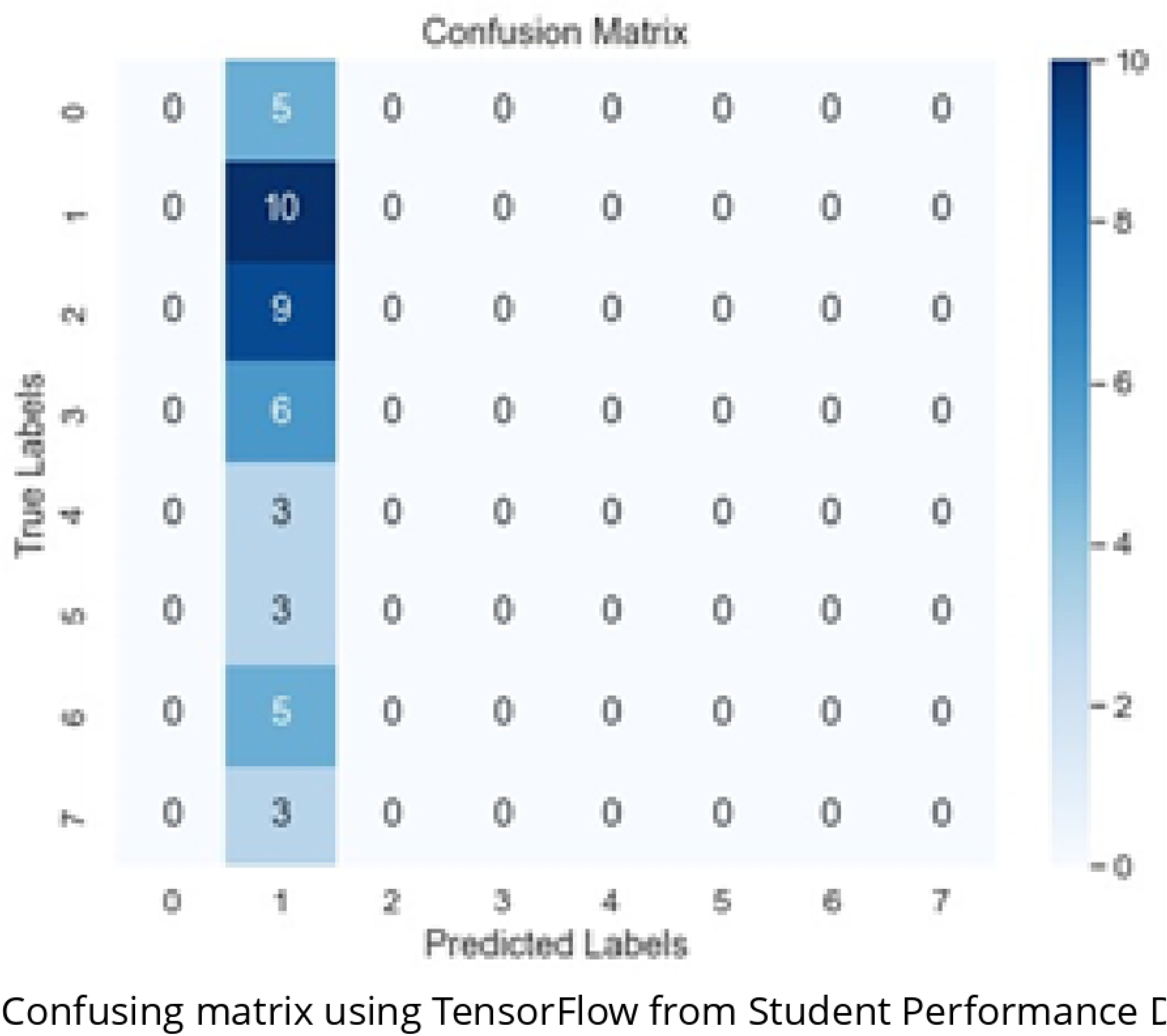
Confusing matrix using TensorFlow from Student Performance Dataset.

Fig 9 depicts a confusion matrix with all diagonal elements equal to zero. This indicates that the model did not correctly predict any instances for any class. This suggests that the model struggles to classify instances across all eight classes. The value in the row labeled “0” and the column labeled “1” (element [0, 1]) is 5. This means that the model incorrectly predicted that five instances that belong to class 1 belong to class 0. Similarly, the value in the row labeled “1” and the column labeled “1” (element [1, 1]) is 10. This indicates that the model incorrectly predicted that ten instances that belong to class 1 belong to class 1. These values suggest that the model cannot distinguish between the different classes. This is likely due to several factors, such as the classification task’s complexity, the training data’s quality, or the model architecture’s limitations.

Fig 10 represents the results of a classification model’s predictions on a multi-class problem with eight different classes using Pytorch from the Student Performance Dataset in Table 6.

**Fig 10.**
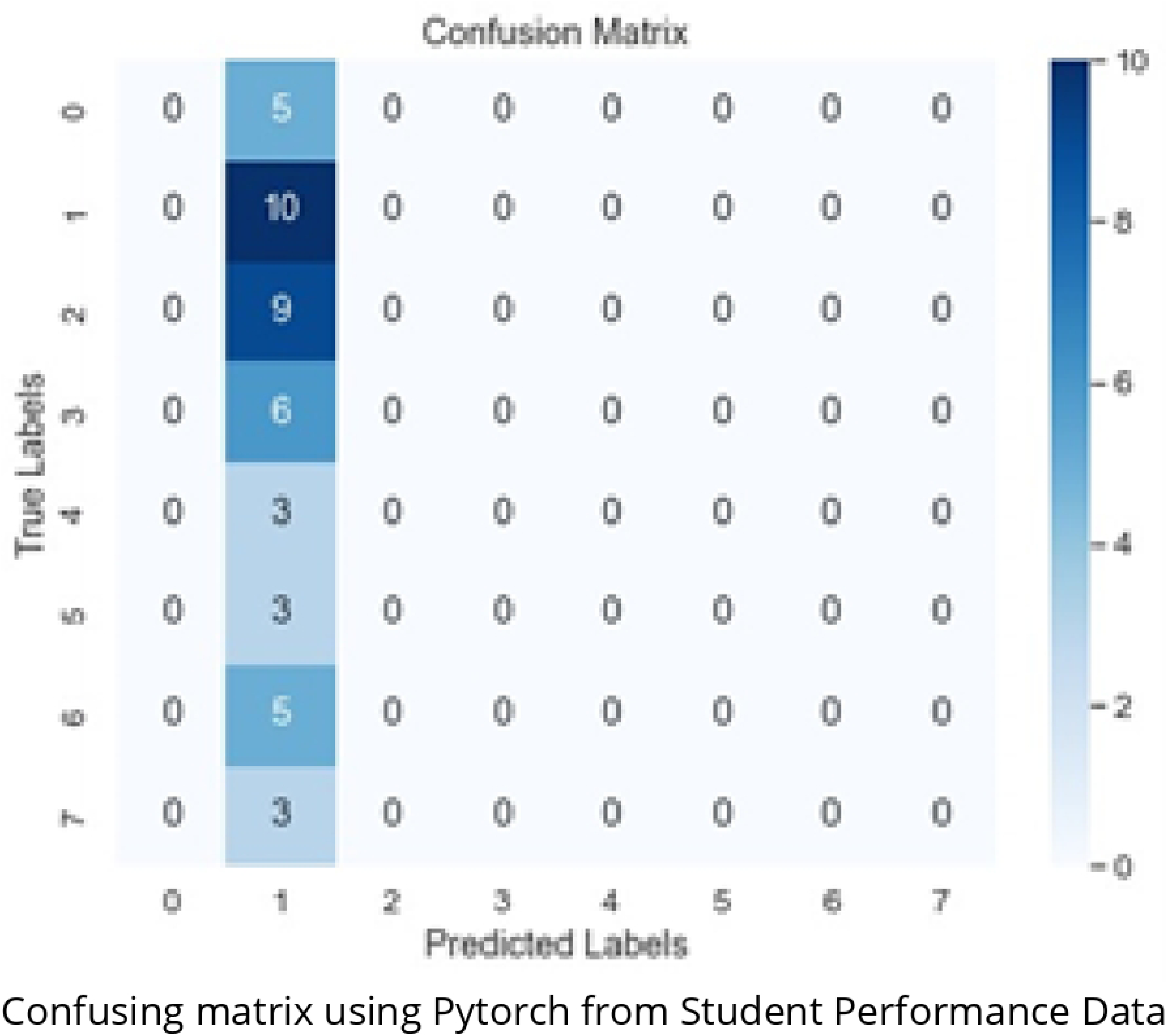
Confusing matrix using Pytorch from Student Performance Dataset.

Fig 10 shows a confusion matrix with identical results obtained from TensorFlow and PyTorch machine learning libraries. This suggests that the two libraries provide consistent performance assessments when applied to the same dataset and model. The confusion matrix provides a detailed breakdown of the model’s predictions, showing the number of instances correctly or incorrectly classified for each class. The matching results suggest that the implementations in both libraries produce equivalent outcomes, making them reliable choices for evaluating and deploying machine learning models. This consistency is essential for ensuring the reliability and reproducibility of model evaluations across different frameworks and environments.

## Discussion

This study systematically benchmarks three prominent machine learning libraries: PyTorch, Keras with TensorFlow backend, and Scikit-learn. The evaluation encompasses two diverse datasets, “College Attending Plan Classification” and “Student Performance,” under standardized criteria, software, and hardware settings. Here, we provide an in-depth discussion of the findings and their implications. Performance on the “College Attending Plan Classification” Dataset The “College Attending Plan Classification” dataset results reveal a substantial performance gap between deep learning frameworks, PyTorch and Keras with TensorFlow backend, and the traditional machine learning library, Scikit-learn. PyTorch and Keras with TensorFlow backend demonstrate exceptional precision, Recall, and F1-score for both classes, showcasing their prowess in handling complex classification tasks. This performance gap can be attributed to the inherent capabilities of deep learning frameworks in capturing intricate patterns. Scikit-learn, while exhibiting commendable performance on this dataset, falls behind its deep-learning counterparts. It suggests that deep learning frameworks are better equipped to handle tasks demanding high predictive accuracy and intricate pattern recognition. PyTorch’s dynamic computation graph and TensorFlow’s computational power give them a significant advantage in these scenarios. Performance on the “Student Performance” Dataset, Conversely, on the “Student Performance” dataset, the three libraries—PyTorch, Keras with TensorFlow backend, and Scikit-learn—deliver more comparable results. While PyTorch and Keras with TensorFlow backend outperform Scikit-learn slightly, all three libraries achieve similar levels of accuracy. This indicates that the complexity of this dataset does not significantly favor any single library. However, it’s noteworthy that Scikit-learn performs the least effectively on this dataset, implying that it might not be the best choice for tasks that require high predictive accuracy in this specific context. Implications and Considerations: the findings underscore the significance of selecting an appropriate machine-learning library based on project requirements. PyTorch and Keras with TensorFlow backend emerge as the top choices for complex classification tasks demanding high predictive accuracy and intricate pattern recognition. These deep learning frameworks exhibit the capability to excel in such scenarios. However, it’s crucial to consider the nature of the dataset and the computational resources available. Scikit-learn remains a valuable and user-friendly option for less complex tasks or resource-constrained environments. This suggests that the choice of the library should not solely hinge on performance metrics but also factors like project complexity and available resources.

## Conclusion

In conclusion, this benchmarking study significantly contributes to machine learning library selection and performance evaluation. Through a systematic and in-depth comparison of three leading libraries—PyTorch, Keras with TensorFlow backend, and Scikit-learn—across two diverse datasets, “College Attending Plan Classification” and “Student Performance,” utilizing feedforward neural networks (FNNs), we have gained valuable insights into the strengths and weaknesses of each library.

This comprehensive analysis provides a clear understanding of the performance variations among these libraries under controlled conditions. It empowers machine learning practitioners to make informed decisions when selecting a library for their project requirements. Importantly, we emphasize that the library choice should not be solely based on performance metrics but also consider factors such as project complexity and available computational resources. Furthermore, this study bridges the gap between theoretical considerations and practical implementation by aligning library performance with real-world datasets and scenarios. It offers pragmatic insights for practitioners navigating the complexities of machine learning, ensuring they leverage the most suitable tools and methodologies for their projects. Comparative studies like this are essential in advancing machine learning practices as the machine learning landscape continues to evolve. By equipping practitioners with knowledge and guidance, we contribute to improving machine learning projects fostering innovation and efficiency in the field. Ultimately, our goal is to empower practitioners to make well-informed decisions, enhancing the quality and impact of their machine-learning endeavors.

## Future work

This benchmarking study provides a valuable foundation for future machine learning library evaluation work. Here are some specific directions that could be explored:

- Expand the evaluation to include emerging libraries and frameworks. This would ensure that the benchmarking results reflect the ever-evolving landscape of machine learning tools.
- Analyze library performance on dynamic datasets of varying domains and complexities. This would provide a more nuanced understanding of how libraries adapt to different data characteristics, which is essential for their practical applicability.
- Evaluate the scalability of libraries for large datasets and distributed computing environments. This is important for libraries intended to be used in production settings.
- Incorporate user experience metrics into the evaluation. This would include ease of use, documentation quality, and community support. These aspects play a vital role in practically adopting machine learning libraries.

By pursuing these and other directions, future work can further advance the understanding of machine learning library performance and usability. This would ultimately benefit machine learning practitioners by providing the tools and information to make informed decisions about library selection.

## Supporting information

**Table 7.**
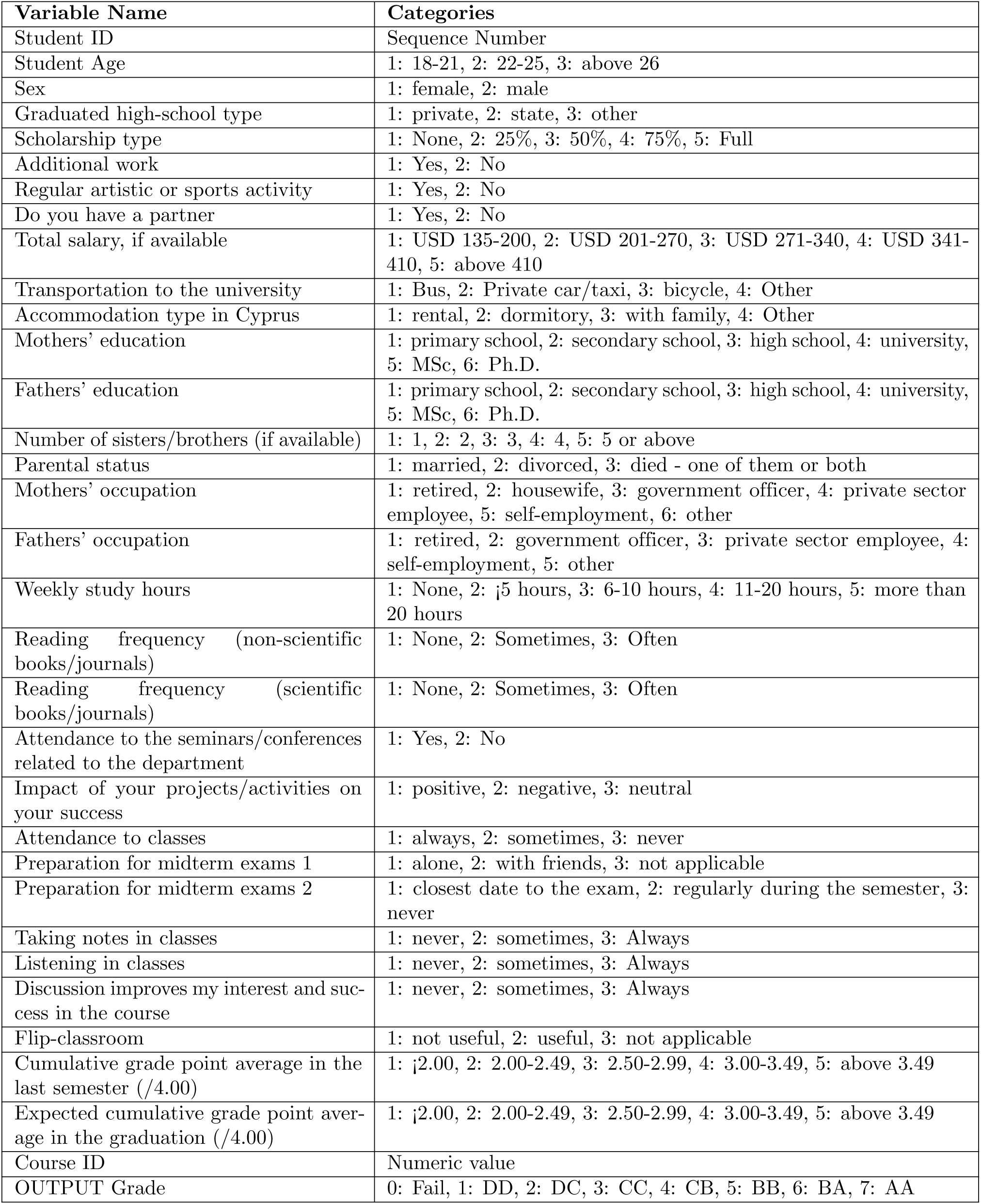
Student Performance Dataset Variables.

**Table 8.** College Attending Plan Classification Dataset Variables.

| Variable | Explanation | Range |
| --- | --- | --- |
| Student ID | The unique identification number of the student | 1, 2, ..., 8000 |
| Gender | The gender of the student | {male (0), female (1)} |
| Parent income | The annual income of the parents (in US dollars) | [4500, 82390] |
| IQ | The IQ of the student in the last test | [60, 140] |
| Encourage | The parents encourage their child to go to college | {encourage, not encourage} |
| Plan | The student eventually plans to go to college | {plan, not plan} |

